# Generation of human neural retina transcriptome atlas by single cell RNA sequencing

**DOI:** 10.1101/425223

**Authors:** Samuel W. Lukowski, Camden Y. Lo, Alexei Sharov, Quan H. Nguyen, Lyujie Fang, Sandy S.C. Hung, Ling Zhu, Ting Zhang, Tu Nguyen, Anne Senabouth, Jafar S. Jabbari, Emily Welby, Jane C. Sowden, Hayley S. Waugh, Adrienne Mackey, Graeme Pollock, Trevor D. Lamb, Peng-Yuan Wang, Alex W. Hewitt, Mark Gillies, Joseph E. Powell, Raymond C.B. Wong

## Abstract

The retina is a highly specialized neural tissue that senses light and initiates image processing. Although the functional organisation of specific cells within the retina has been well-studied, the molecular profile of many cell types remains unclear in humans. To comprehensively profile cell types in the human retina, we performed single cell RNA-sequencing on 20,009 cells obtained post-mortem from three donors and compiled a reference transcriptome atlas. Using unsupervised clustering analysis, we identified 18 transcriptionally distinct clusters representing all known retinal cells: rod photoreceptors, cone photoreceptors, Müller glia cells, bipolar cells, amacrine cells, retinal ganglion cells, horizontal cells, retinal astrocytes and microglia. Notably, our data captured molecular profiles for healthy and early degenerating rod photoreceptors, and revealed a novel role of *MALAT1* in putative rod degeneration. We also demonstrated the use of this retina transcriptome atlas to benchmark pluripotent stem cell-derived cone photoreceptors and an adult Müller glia cell line. This work provides an important reference with unprecedented insights into the transcriptional landscape of human retinal cells, which is fundamental to our understanding of retinal biology and disease.

## Introduction

The eye is a highly specialised sensory organ in the human body. Sight is initiated by the conversion of light into an electrical signal in the photoreceptors of the neurosensory retina. The rod photoreceptors are responsible for light detection at extremely low luminance, while the cone photoreceptors are responsible for colour detection and operate at moderate and higher levels. Following preprocessing, by horizontal, bipolar and amacrine cells, the resultant signal is transferred via ganglion cells to the brain. Neurotransmitter support is provided by Müller glia, retinal astrocytes and microglial cells. Inherited retinal diseases are becoming the leading cause of blindness in working age adults, with loci in over 200 genes associated with retinal diseases (RetNet: https://sph.uth.edu/retnet/), often involving specific retinal cell types. Knowledge of the transcriptome profile of individual retinal cell types in humans is important to understand the cellular diversity in the retina, as well as the study of retinal genes that contribute to disease in individual retinal cell types. (Hornan *et al*, 2007a; Mustafi *et al*, 2016a; Whitmore *et al*, 2014a; Farkas *et al*, 2013a; Pinelli *et al*, 2016a)

The transcriptome profiles of whole human retina from adults (Hornan *et al*, 2007b; Mustafi *et al*, 2016b; Whitmore *et al*, 2014b; Farkas *et al*, 2013b; Pinelli *et al*, 2016b) and during fetal development (Hoshino *et al*, 2017; Kozulin *et al*, 2009) have been previously described. However, these studies only assayed the averaged transcriptional signatures across all cell types, meaning that the cellular heterogeneity in the retina is lost. As such, the transcriptional pathways that underlie the highly specialised function of many human retinal cell types remain unclear; including the rod and cone photoreceptors, Müller glia cells, horizontal cells, and amacrine cells. Recent advances in RNA sequencing and microfluidic platforms have dramatically improved the accessibility of single cell transcriptomics, with increased throughput at a lower cost. Critically, single-cell microfluidics and low-abundance RNA library chemistries allow accurate profiling of the transcriptome of individual cell types. This has been demonstrated in the mouse, where transcriptome profiles of the mouse retina (Macosko *et al*, 2015) and retinal bipolar cells (Shekhar *et al*, 2016) have been described at the single cell level using the Drop-seq method (Macosko *et al*, 2015). These studies provided a molecular classification of the mouse retina and identified novel markers for specific cell types, as well as novel candidate cell types in the retina. Recently, single cell transcriptomics was used to analyse the human retina. Phillips et al. have profiled a total of 139 adult retina cells using the C1 Fluidigm platform (Phillips *et al*, 2018), but the limited number of profiled cells presents challenges in the annotation and accurate identification of individual retinal cell types. Moreover, a flow cytometry approach was used to isolate 65 human fetal cone photoreceptors followed by scRNA-seq profiling (Welby *et al*, 2017).

Herein we report the generation of a human neural retina transcriptome atlas using 20,009 single cells from three donors. Our data provide new insights into the transcriptome profile of major human retinal cell types and establish a high cellular-resolution reference of the human neural retina, which will have implications for identification of biomarkers and understanding retinal cell biology and diseases.

## Results

### Preparation of human neural retinal samples and generation of single cell transcriptome atlas

We obtained post-mortem human adult eyes approved for research purposes following corneal transplantation. As the transcriptome profile of human retina pigment epithelium cells has already been reported (Strunnikova *et al*, 2010; Liao *et al*, 2010), we focused solely on the neural retina layers. In this study we extracted the neural retina from twelve donor eyes (Supplementary Table 1). We observed consistent cell viability across retinal tissues retrieved within 15 hours post-mortem (Supplementary figure 1A) and found that donor age does not impact negatively on cell viability in the extracted neural retina (Supplementary figure 1B). To minimize potential risk of mRNA degradation due to reduced cell viability, we selected three donor samples retrieved within 15 hours post-mortem and analysed them with single cell RNA sequencing (scRNA-seq) using the 10X Genomics Chromium platform.

Sequence data from five scRNA-seq libraries derived from the three neural retinal samples were pooled for processing and analysis. From 23,000 cells, we obtained an average of 40,232 reads per cell and 1,665 UMIs (unique transcripts) per cell. Following quality control and filtering in Seurat, our final dataset contained 20,009 cells, which were taken forward for further analysis.

The scRNA-seq data was initially analysed using an unsupervised graph clustering approach implemented in Seurat (version 2.2.1) to classify individual cells into cell subpopulations according to similarities in their transcriptome profiles. Overall, the cells were classified into 18 transcriptionally distinct clusters (Figure 1A, Supplementary figure 2). We first assessed the variation between donor samples and between library preparations (Supplementary table 2, supplementary figure 3 and 4). Interestingly, although many of the identified clusters are present in all three donor retinal samples, we also observed several donor-specific clusters corresponding to rod photoreceptors (Figure 1B, Supplementary figure 3A and 3B). In contrast, we observed minimal variation between two different libraries prepared from the same donor sample, supporting the quality of the scRNA-seq datasets in this study (Supplementary figure 4).

**Figure 1:**
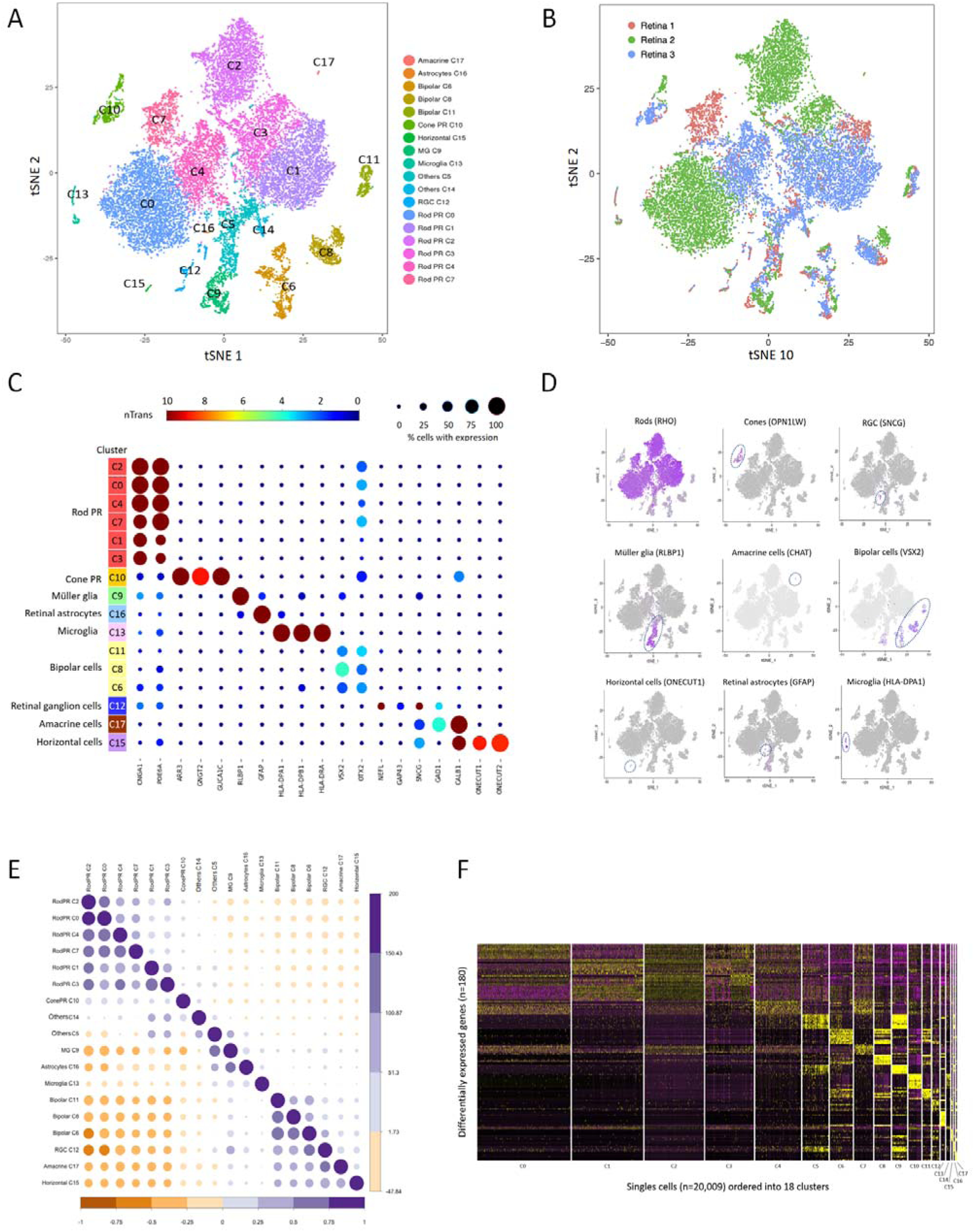
Single cell transcriptome atlas for human neural retina. t-SNE visualization of 20,009 human retinal cells colored by A) annotation of 18 transcriptionally distinct clusters (C0-C17) and B) their distribution in 3 donor retina samples. C) Feature expression heatmap showing expression patterns of major retinal class markers across 16 retinal cell clusters. The size of each circle depicts the percentage of cells expressing the marker within the cluster. Brown colour indicates ≥10 nTrans (number of transcripts). D) t-SNE plots showing expression of a set of selected marker genes for major retinal classes. E) Correlation matrix for the identified 18 clusters. The upper triangle depicts the z-value for correlation and the lower triangle depicts the correlation coefficient for gene expression in clusters. F) Heatmap of differentially expressed genes used to classify cell types for each cluster compared to all other clusters for the 18 retinal cell clusters. The rows correspond to top 10 genes most selectively upregulated in individual clusters (p< 0.01, Benjamini-Hochberg correction) and the columns show individual cells ordered by cluster (C0-C17).

**Figure 2:**
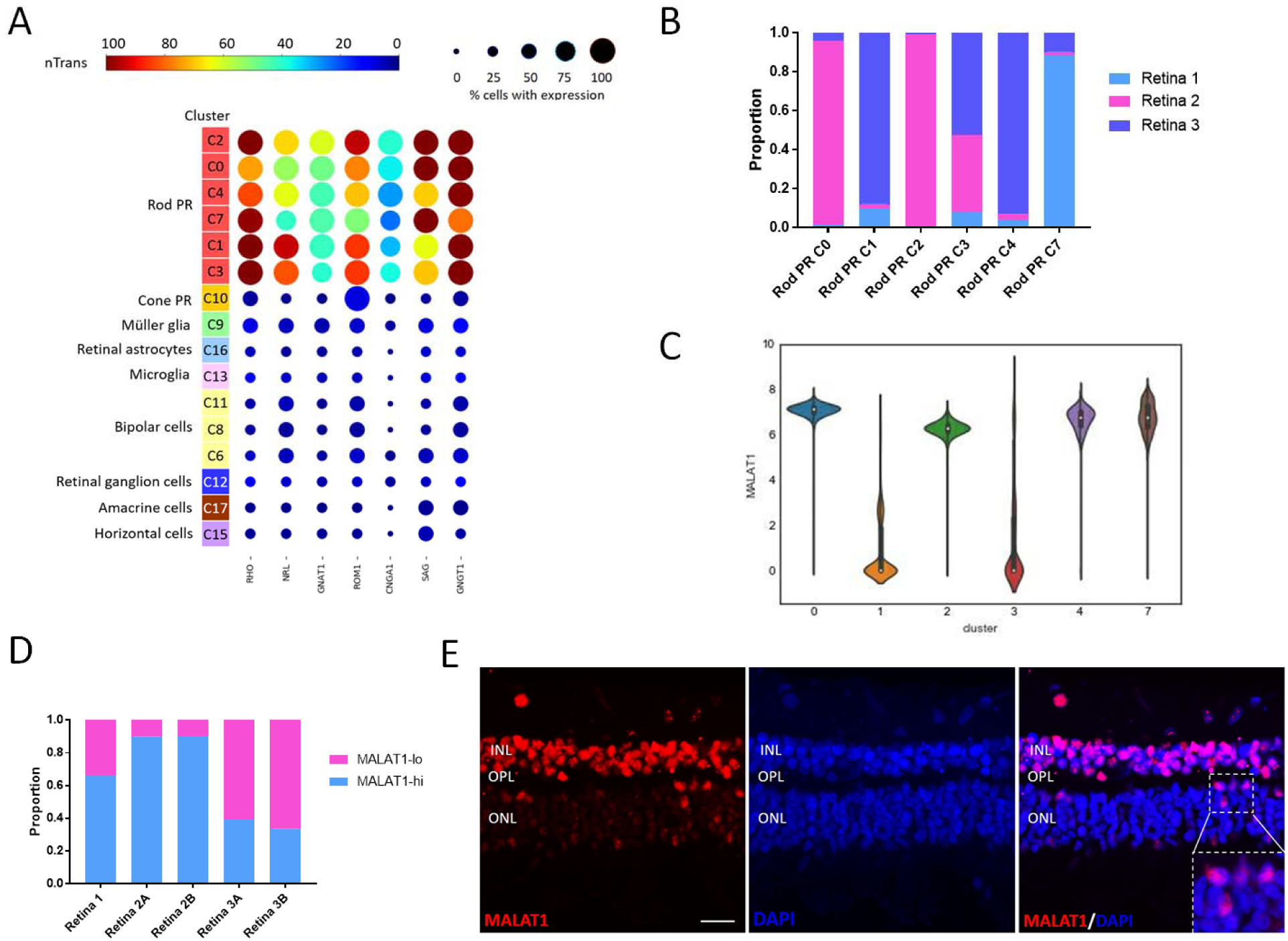
Identification of *MALAT1-hi* and *MALAT1-lo* subpopulations of rod photoreceptors. A) Feature expression heatmap of a panel of known marker genes for rod photoreceptors across the identified 16 retinal cell clusters. Brown colour indicates ≥100 nTrans (number of transcripts). B) Representation of the three donor retina samples in the six rod photoreceptor clusters. C) Violin plot showing high or low expression levels of *MALAT1* in rod photoreceptor clusters. D) Distribution of rod photoreceptor populations with high MALAT1 expression (*MALAT1*-hi) or low *MALAT1* expression (*MALAT1*-lo) in three donor retin samples. E) Fluorescent in situ hybridization analysis of human peripheral retina showing heterogeneous levels of *MALAT1* expression in the rod photoreceptors located in the outer nuclear layer (ONL). INL: inner nuclear layer; OPL: outer plexiform layer. Scale bar = 20µm.

**Figure 3:**
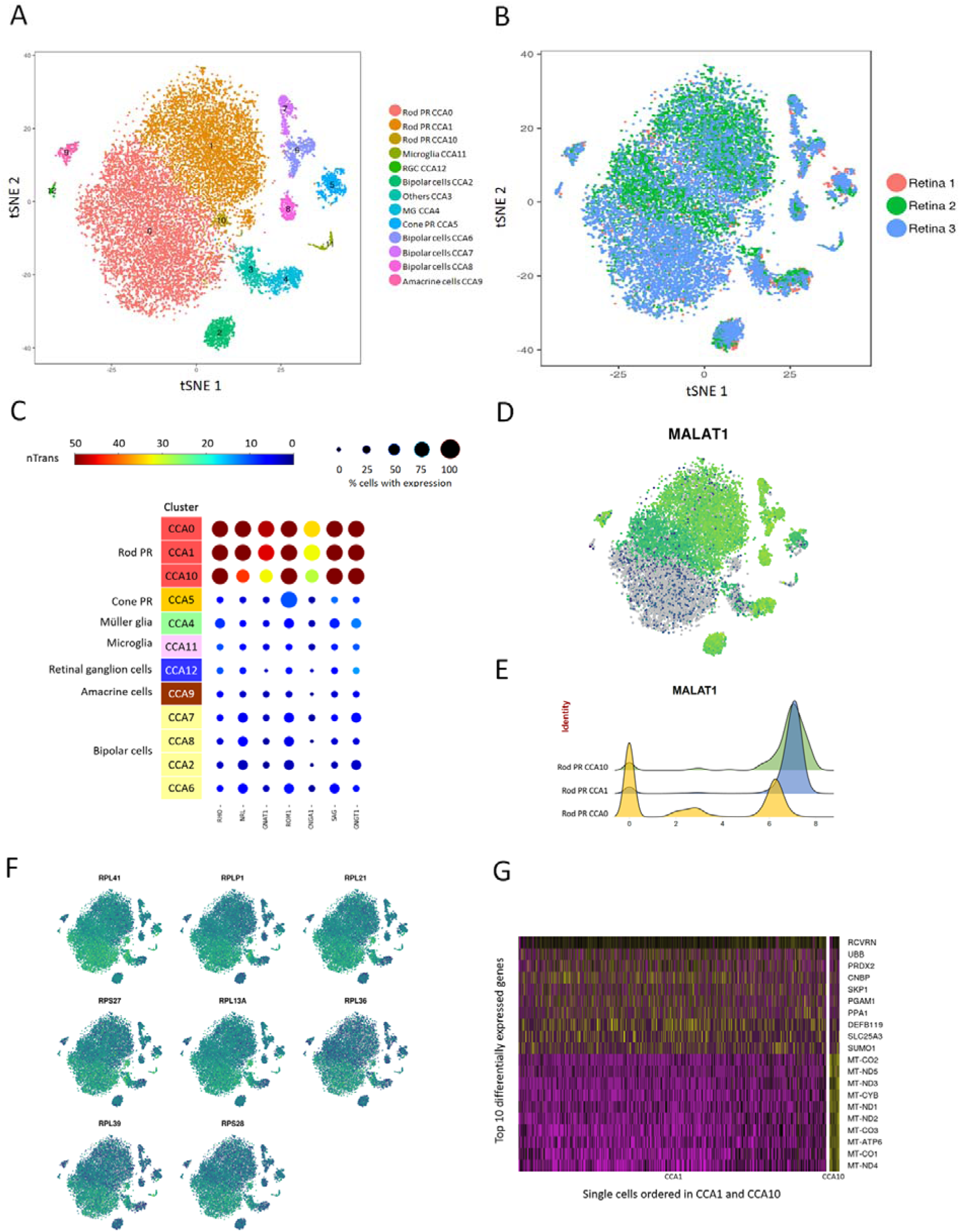
***MALAT1* subpopulations of rod photoreceptors is not due to donor variation.** Canonical correlation analysis was used to effectively correct donor-specific variations in rod photoreceptors. A) t-SNE visualization of human retinal cells colored by annotation of 13 transcriptionally distinct clusters (CCA0-CCA12) and B) their distribution in 3 donor retina samples. C) Feature expression heatmap showing expression patterns of 7 rod photoreceptor markers across 12 retinal cell clusters. The size of each circle depicts the percentage of cells expressing the marker within the cluster. Brown colour indicates ≥50 nTrans (number of transcripts). D) t-SNE plots showing expression of *MALAT1*. E) Expression pattern of *MALAT1* in the rod photoreceptor showing *MALAT1-hi* (CCA1, CCA10) and *MALAT1-lo* (CCA0) subpopulations. x-axis depicts normalized transcript levels. F) t-SNE plots showing expression of major ribosomal genes. G) Heatmap of differentially expressed genes between the two *MALAT1-hi* clusters CCA1 and CCA10. The rows correspond to top 10 genes most selectively upregulated in individual clusters (p< 0.01, Benjamini-Hochberg correction) and the columns show individual cells ordered in CCA1 and CCA10.

**Figure 4:**
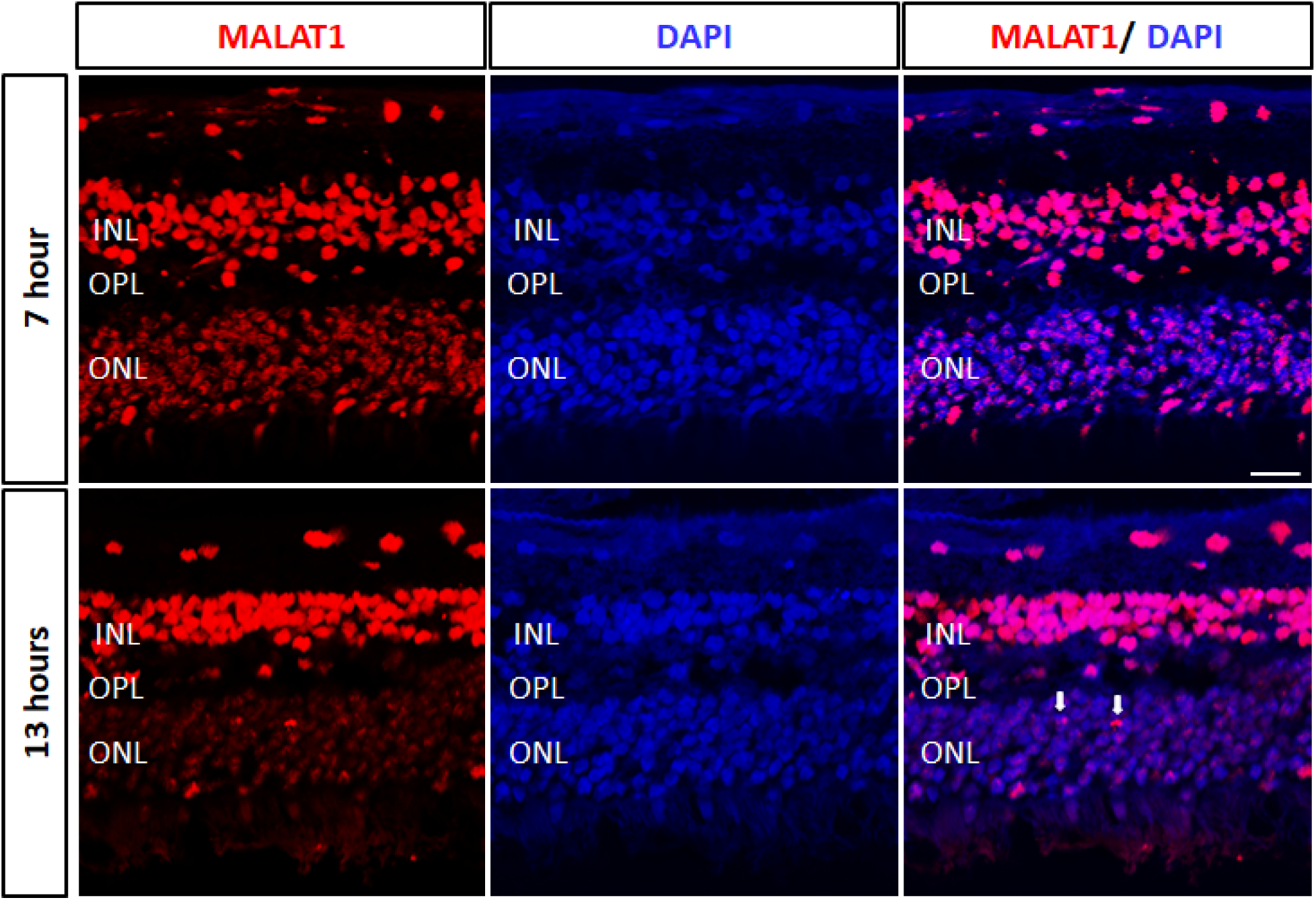
Loss of *MALAT1* expression in rod photoreceptors following post-morte degeneration. Fluorescent in situ hybridization analysis of the same donor human peripheral retina at different time points post-mortem (7 and 13 hours, Retina 7), showing decreases in *MALAT1-hi* rod subpopulations in the outer nuclear layer (ONL) at later time point. INL: inner nuclear layer; OPL: outer plexiform layer. Scale bar = 20µm. White arrows indicated *MALAT1-hi* rod photoreceptors.

### Identification of major cell types in the human retina using scRNA-seq

Based on known markers (Klimova *et al*, 2015; Vecino *et al*, 2016; Shekhar *et al*, 2016; Macosko *et al*, 2015; Soto *et al*, 2008; Imanishi *et al*, 2002; Blackshaw *et al*, 2001; Corbo *et al*, 2007), we were able to assign cell identities to the 16 of the 18 clusters (Figure 1A, 1C, 1D), corresponding to rod photoreceptors (*PDE6A, CNGA1, RHO*), cone photoreceptors (*ARR3, GNGT2, GUCA1C*), Müller glia (*RLBP1/CRALBP*), retinal astrocytes (*GFAP*), microglia (*HLA-DPA1, HLA-DPB1, HLA-DRA*), bipolar cells (*VSX2, OTX2*), retinal ganglion cells (*NEFL, GAP43, SNCG*), amacrine cells (*GAD1, CALB1, CHAT*) and horizontal cells (*ONECUT1, ONECUT2*). The expression of selected marker genes are displayed in *t*-SNE plots (Figure 1D). Two clusters (C5 and C14) express markers from multiple retinal cell types (Supplementary figure 5), thus we were unable to assign cell identities to these 2 clusters and they were excluded from further analysis. Interestingly, our data demonstrated multiple transcriptionally distinct clusters within the rod photoreceptors (6 clusters) and bipolar cells (3 clusters). In contrast, only one cluster was detected for cone photoreceptors, Müller glia, retinal ganglion cells, horizontal cells, amacrine cells, retinal astrocytes and microglia respectively. Correlation analysis confirmed the similarity between clusters within the same cell type (Figure 1E). As expected, we observed high correlations between the expression levels of transcripts within photoreceptor cell types (rod and cones), as well as glial cells (retinal astrocytes and Müller glia) and other retinal neurons (bipolar cells, retinal ganglion cells, amacrine cells and horizontal cells). The composition of cell populations across our three donors show that the majority of the cells in human neural retina were rod photoreceptors (∼74%) followed by bipolar cells (∼10%) . These results are similar to those reported in mice, where rod photoreceptors and bipolar cells form the majority of cells in the retina (Jeon *et al*, 1998; Macosko *et al*, 2015).

**Figure 5:**
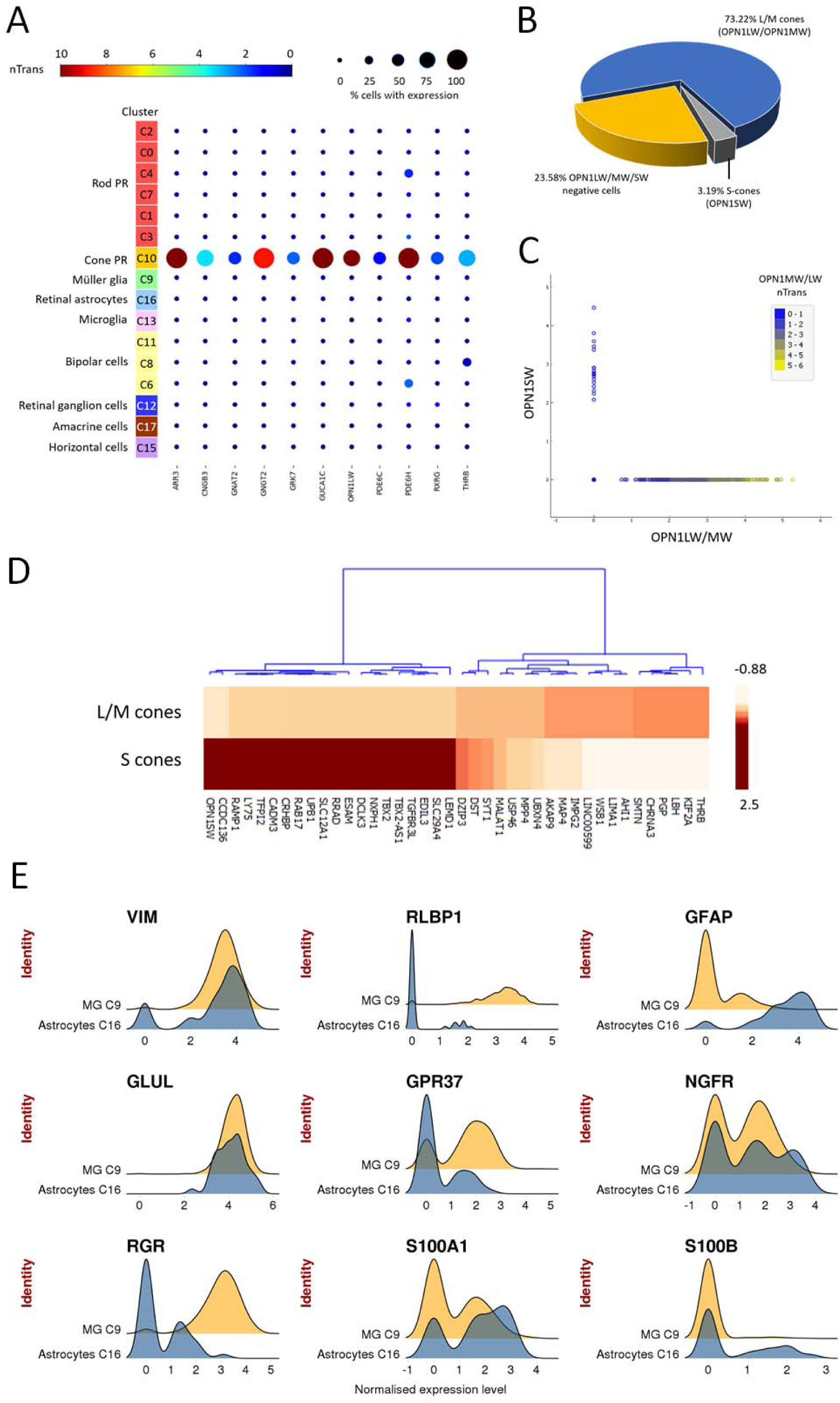
Assessment of cone photoreceptor and glial cell types in human retina . A) Feature expression heatmap showing the expression of 11 known cone photoreceptor markers across 16 retinal cell clusters. Brown colour indicates ≥10 nTrans (number of transcripts). B) The proportion of cone photoreceptor subtypes identified in C10 cluster, based on expression of *OPN1LW / OPN1MW* (L/M cones) and *OPN1SW* (S cones). C) Scatter plots showing expression of *OPN1LW/OPN1MW* or *OPN1SW* in individual cone photoreceptors for C10 cluster. The colour depicts expression level for *OPN1LW/OPN1MW* in individual cells. D) Heatmap of top 20 differentially expressed genes between L/M cones and S cones. The colour depicts normalised gene expression (z-score capped at 2.5). E) Expression pattern of glial markers in Muller glia (C9) and retinal astrocytes (C16). x-axis depicts normalized transcript levels.

To identify genes whose expression was specific to a given cell type, we performed differential gene expression analysis to identify marker genes for each cluster (Figure 1F). We subsequently extracted membrane-related proteins from gene ontology annotations to identify surface markers, which can be used to develop immuno-based methods to isolate primary culture of individual retinal cell types. Supplementary table 3 lists the identified surface markers for individual retinal cell types. We also assessed the gene expression of a panel of commonly known markers in amacrine cells and bipolar cells (Supplementary figure 6-7), as well as a panel of markers for subtype identification recently identified in mouse scRNA-seq studies (Shekhar *et al*, 2016; Macosko *et al*, 2015). In summary, we profiled the transcriptomes of all major cell types in the human retina in the presented dataset. Due to their abundance, for the subsequent analyses we focused on the photoreceptors and glial cells.

### Profiling healthy and degenerating human rod photoreceptor subpopulations

We profiled 14,759 rod photoreceptors and showed that they can be classified into six populations with distinct gene expressions (c0, c1, c2, c3, c4, c7; Supplementary table 3). We assessed these six clusters with a panel of 7 known rod or pan-photoreceptor markers (Figure 2A). Our results suggest differential expression patterns among the 7 markers. All 7 rod markers are highly abundant, consistent with previous scRNA-seq studies of mouse and human retina (Phillips *et al*, 2018; Macosko *et al*, 2015). The 7 markers showed differential expression patterns in the 6 identified rod photoreceptor clusters. In particular, *RHO, GNGT1* and *SAG* have the highest levels of rod marker detected, followed by *NRL*, *ROM1*, *GNAT1* and *CNGA1.* We also noted that *ROM1* is expressed in both rod and cone photoreceptors, which is consistent with previous studies (Boon *et al*, 2008). Importantly, many rod photoreceptor clusters consist of a majority of cells from a single donor (>90% for c0, c2, c4 and >80% for c1, c7; Figure 2B). It is possible that this observation is due to the systematic biases such as differences in tissue retrieval time, age of donors, or other sample preparation variation. The exception is cluster c3, which is well represented by all three donors.

Next we set out to further define and classify heterogeneity in rod photoreceptors. We observed that *MALAT1*, a long non-coding RNA that plays a role in retinal homeostasis and disease (Wan *et al*, 2017), was robustly expressed in ∼66% of the identified rod photoreceptors (9,750 cells) while the rest had little to no expression (5,009 cells; Figure 2C). As such, we utilized *MALAT1* expression as a discriminator and investigated differences between rod photoreceptors with high expression (*MALAT1*-*hi*; > 4.68 normalised transcripts per cell) or low expression (*MALAT1-lo*; < 4.68 normalised transcripts per cell). *MALAT1*-*hi* and *-lo* rod photoreceptors were consistently found in each donor and library samples, with *MALAT1*-*hi* accounting for ∼66%, 90% and 36% of the rods in donors #1, #2 and #3 respectively (Figure 2D). To further validate this finding, we performed RNA *in situ* hybridization in another three donor retinal samples. We consistently observed the presence of *MALAT1-hi* and *-lo* subpopulations of rod photoreceptors in all retinal samples (Figure 2E, Supplementary figure 8). Together, these results showed the presence of heterogeneity within rod photoreceptors that can be distinguished by *MALAT1* expression.

To rule out the possibility that the presence of *MALAT1* rod subpopulations is due to donor sample variations, we applied Canonical Correlation Analysis (CCA) to the dataset, via the Seurat package, to correct for technical and batch artefacts. We found that CCA can effectively corrected the donor-specific effect on rod photoreceptor clusters (Figure 3A and 3B). Following CCA correction, we identified 3 rod photoreceptors clusters (CCA0, CCA1, CCA10), which expressed a panel of 7 rod photoreceptor markers and were well represented in all donor samples (Figure 3C). Notably, the majority of cells in CCA0 were low in *MALAT1* expression, while CCA1 and CCA10 represented *MALAT1-high* rod subpopulations (Figure 3D and 3E). This is consistent with our RNA *in situ* hybridization analysis, where we consistently observed *MALAT1-hi* and *-lo* subpopulations of rod photoreceptors in all retinal samples (Supplementary figure 9). Collectively these results supported that the *MALAT1* heterogenity in rod photoreceptors are not due to inter-individual variability.

We also considered the possibility that *MALAT1-lo* rod subpopulations may represent an artefact of ‘low quality cells’ in scRNA-seq data, due to a low number of sequencing reads or broken cell membrane. In this regard, up-regulated levels of mitochondrial-encoded genes and ribosomal proteins can be used to identify such low quality cells in scRNA-seq data (Ilicic *et al*, 2016). For our scRNA-seq dataset, we did not observe upregulation in gene expression for a panel of ribosomal proteins (*RPL41, RPLP1, RPL21, RPS27, RPL13A, RPL36, RPL39* and *RPS28*; Figure 3F). However, the rod cluster CCA10, representing 1.4% of rod photoreceptor cells, showed markedly increased levels of mitochondrial-encoded genes (*MT-CO2, MT-ND5, MT-ND3, MT-CYB, MT-ND1, MT-ND2, MT-CO3, MT-ATP6, MT-CO1, MT-ND4)*, suggesting that CCA10 represented a low-quality *MALAT1-hi* rod cluster and was excluded from further analysis.

As we utilised post-mortem retinal samples in this study, we reasoned that *MALAT1-lo* subpopulation may reflect the early stages of post-mortem degeneration in rod photoreceptors. To determine this, we extracted retinal samples from the same donor at different time points of progressive post-mortem degeneration, with longer time points predicted to have more stressed/dying photoreceptors. Our results showed that there was a high proportion of *MALAT1-hi* rod photoreceptors at 7 hours post-mortem (Figure 4, ∼95%). However, we observed a marked decrease in *MALAT1* expression in rod photoreceptors at 13 hours post-mortem. Similar results were observed for the three retinal samples processed for scRNA-seq (Supplementary figure 8B). Together, these results demonstrated that *MALAT1* is a novel marker for healthy photoreceptors with loss of expression preceding putative cell degeneration. In summary, we showed that scRNA-seq can be used to profile healthy (CCA1) and degenerating rod photoreceptors (CCA0), which can be distinguished by high or low *MALAT1* expression levels respectively.

### Transcriptome profile of cone subtypes in the human retina

We detected 564 cone photoreceptor cells in our scRNA-seq data, which are distinguishable from the other cell types by the expression of the cone marker genes *ARR3, CNBG3, GNAT2, GNGT2, GRK7, GUCA1C, PDE6C, PDE6H, OPN1LW, RXRG* and *THRB* (Figure 5A). All 11 marker genes analysed show specific expression patterns in the cone cluster (C10). We set out to further assess the composition of the cone cluster. In humans, there are three identified subtypes of cone photoreceptor which can be distinguished by expression of a sole opsin gene: *OPN1SW*-positive S-cones, *OPN1MW*-positive M-cones and *OPN1LW*-positive L-cones respond preferentially to shorter, medium and longer wavelengths responsible for colour vision (Roorda & Williams, 1999). Notably *OPN1LW* and *OPN1MW* exhibit ∼98% sequence homology and are unable to be distinguished by 3’ sequencing utilised in this study. By quantifying the number of cells that express the opsin genes, our results showed that the majority of the cone cluster are L/M cones (73.22%) and S-cones in much lower proportion (3.19%, Figure 5B), at levels consistent with those estimated by a previous adaptive optics and photo-bleaching study (Roorda & Williams, 1999). As expected, the identified cone photoreceptors only express either *OPN1SW* or *OPN1LW/MW* (Figure 3C).

To further study the molecular differences and identify molecular markers for the cone subtypes, we performed differential gene expression analysis to determine genes that can distinguish the cone subtypes. Our results identified a panel of genes that differentially marked S-cones (e.g. *CCDC136*, *RAMP1*, *LY75, CADM3, TFPI2, CRHBP, RAB17, UPB1, RRAD, SLC12A1*) and L/M-cones (e.g. *THRB, KIF2A, LBH, PGP, CHRNA3, AHI1, LIMA1*; Figure 3D). Together these results detailed the molecular profiles and identified marker genes that can distinguish the cone subtypes in human.

### Assessment of glial cells in human retina

Next, we focused on two related glial cell types in the human retina, the Müller glia and the retinal astrocytes. Our scRNA-seq data has profiled a total of 2,723 Müller glia cells which classified into a single cluster (C9) and 49 retinal astrocytes which form a single cluster (C16). Figure 5E shows the expression of a panel of 9 commonly used markers for Müller glia and retinal astrocytes. Our results demonstrated that many of these markers are present in both Müller glia and retinal astrocytes at differential expression levels. *VIM, GLUL* and *S100A1* marked both Müller glia and retinal astrocytes at high expression levels. *GFAP* represents a reliable marker for retinal astrocytes, and its expression is consistent with a previous report (Vecino *et al*, 2016). Notably, Müller glia are low in *GFAP* expression, indicating they are not in an activated state commonly caused by stresses and reactive gliosis (Fernández-Sánchez *et al*, 2015). The *S100B* is also expressed in retinal astrocytes at varying levels but absent in Müller glia. Conversely, Müller glia can be distinguished from retinal astrocytes by high expression levels of *RLBP1*, and *RGR* to a lesser extent. Together these results provide insights into the differential expression patterns of known glial markers in Müller glia and retinal astrocytes in human.

As glial cell proliferation has been linked to a range of pathological conditions including retinal gliosis and retinal injury (Subirada *et al*, 2018), this provides a means of assessing the health of the profiled retinas. We assigned a cell cycle phase score to each cell using gene expression signatures for the G1, S, G2, and G2/M phases (Kowalczyk *et al*, 2015); Supplementary figure 9). We determined that most of the Müller glial cells expressed genes indicative of cells in G1 phase (75%), suggesting they are not proliferative. These results demonstrated the absence of hallmarks of gliosis/retinal injury and support the quality of the donor retinas profiled.

### Using the human neural retina transcriptome atlas for benchmarking

To demonstrate the use of our dataset as a benchmarking reference, we compared the scRNA-seq profiles of distinct cell types generated using alternative methods, including fetal human cone photoreceptors, human induced pluripotent stem cell derived-cone photoreceptors (hiPSC-cone; (Welby *et al*, 2017), and a sample of adult human retina with 139 cells (Phillips *et al*, 2018). Correlation analysis demonstrated that the adult human retina sample showed highest similarity to rod photoreceptor (0.63, Supplementary figure 10), which is expected as rod photoreceptors represent the majority of the cells in the retina. Interestingly, our results also showed that the transcriptome of hiPSC-cone after 15 weeks of differentiation exhibited the highest similarity to cone photoreceptors, and low similarities to all other retinal cell types (Figure 6A, Supplementary figure 10). In particular, hiPSC-cone showed high similarities to fetal cone photoreceptors and adult cone photoreceptors (0.71 and 0.61 respectively), and a much lower similarity to adult rod photoreceptors (0.33). In support of this, principal component analysis demonstrated that the hiPSC-cone are closer to fetal cone photoreceptors, rather than the adult counterpart (Figure 6B, Supplementary figure 10). These results confirmed direct differentiation of hiPSCs to cone photoreceptors with good quality, and the hiPSC-derived cone photoreceptors are closer to fetal origin compared to their adult counterpart.

**Figure 6:**
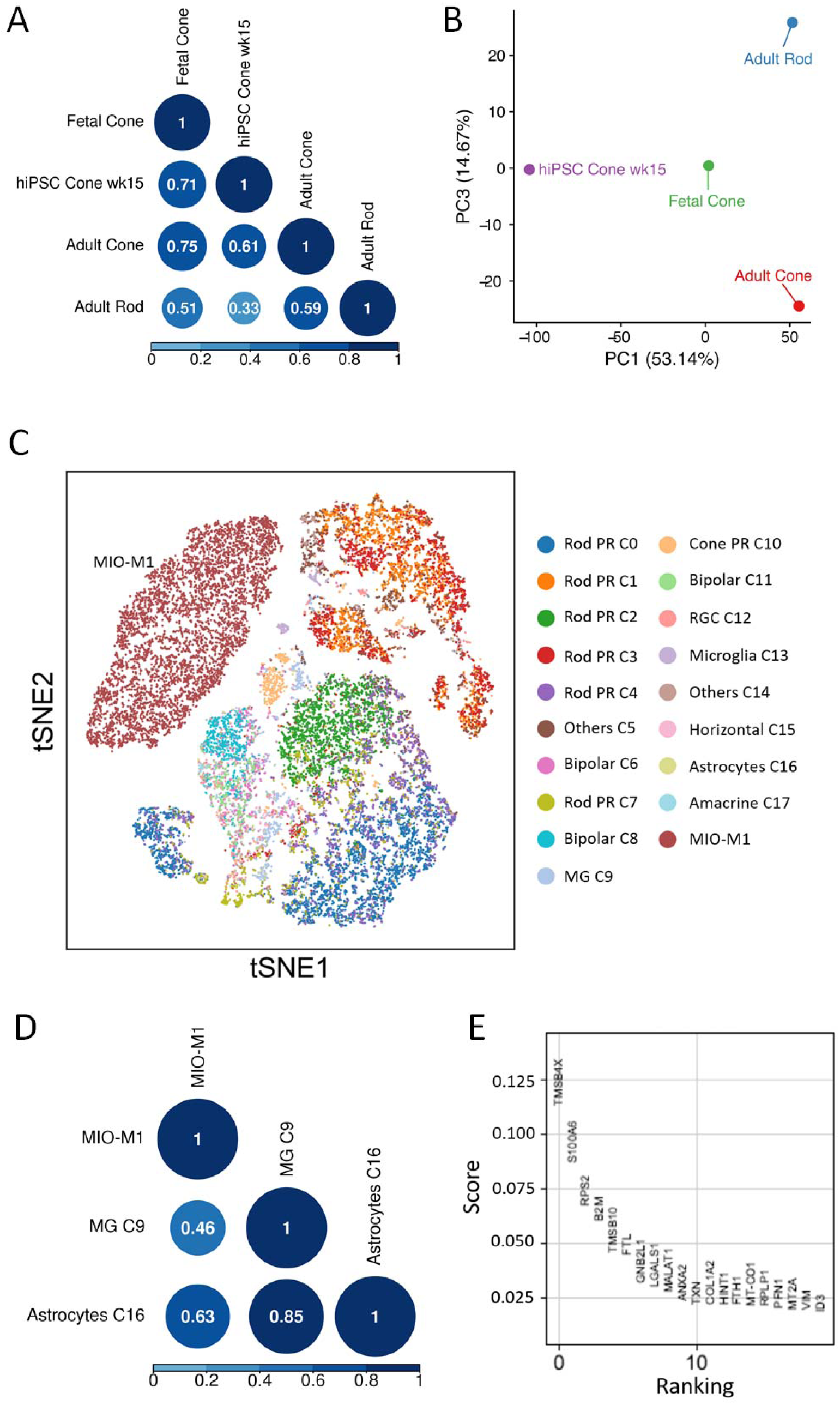
Benchmarking retinal cells using the human neural retina atlas. A) Correlation analysis of scRNA-seq data of hiPSC-derived cone photoreceptors (week 15) against fetal cone photoreceptors (Welby *et al*, 2017), as well as adult cone and rod photoreceptors from this human neural retina atlas. B) Principal component analysis to assess transcriptome similarity of hiPSC-derived cone photoreceptors to fetal and adult photoreceptors. C) t-SNE analysis of the human Müller glia cell line MIO-M1 with the retinal cell types identified in this human neural retina atlas. D) Correlation analysis of MIO-M1 with all major human retinal cell types. E) Top ranked differentially expressed genes identified in MIO-M1 compared to other retinal cell types based on logistic regression score.

In another benchmarking example, we set out to assess the potential differences between *in vitro* cell lines compared to adult cells *in vivo*. In this regard, we compared the spontaneously immortalised human Müller glia cell line MIO-M1 (Lawrence *et al*, 2007; Limb *et al*, 2002) to all the retinal cell types identified in our dataset. Using scRNA-seq, we profiled 7,150 MIO-M1 cells with 23,987 reads per cell post-normalization corresponding to 3,421 detected genes. Unsupervised clustering and t-SNE analysis showed that the MIO-M1 cells formed one cluster that is transcriptionally distinct from all retina cell types identified in the human neural retina dataset (Fig 6C, Supplementary figure 10). Correlation analysis showed that MIO-M1 displayed similarities to retinal glial cells, with higher similarity to astrocytes compared to Müller glia (0.63 and 0.46 respectively, Fig 6D). In particular, we identified that MIO-M1 express high levels of thymosin beta 4 gene (*TMSB4X*), which has been linked to glioma malignancy (Wirsching *et al*, 2013), as well as the calcyclin gene (*S100A6*), which is implicated in macular or cone associated diseases (Yoshida *et al*, 2004); Figure 4E). Together, our results highlighted the similarities and differences of MIO-M1 to adult retinal glial cells in human.

## Discussion

The data presented here describe the generation of a detailed reference transcriptome atlas of the human neural retina at the single-cell level. The establishment of reference transcriptome maps for individual cell types in the retina provide unprecedented insights into the signals that define retinal cell identity and advance our understanding of the retina. This human neural retina transcriptome data can be used as a benchmark to assess the quality and maturity of pluripotent stem cell-derived retinal cells, such as retinal ganglion cells (Gill *et al*, 2016; Kobayashi *et al*, 2018; Sluch *et al*, 2015) and photoreceptors (Lakowski *et al*, 2018). We also confirmed that a relatively low level of sequencing depth (∼40,232 reads per cell) is sufficient for identification and classification of major cell types in a complex tissue like retina. However, we acknowledge that increasing the sequencing depth and cell number could improve the ability to distinguish retinal subtypes that are highly complex, such as the amacrine cells and the retinal ganglion cells.

One of the most interesting observations is the presence of heterogeneous subpopulations within known retinal cell types. This highlights the sensitivity of using a scRNA-seq approach to capture and classify retinal cell types in an unbiased manner. In particular, our results demonstrated the presence of two rod photoreceptor subpopulations in post-mortem retina that display differential expression of *MALAT1*. Notably, the presence of *MALAT1-hi* and -*lo* rod subpopulations were consistently observed in all post-mortem samples analysed (n=7). We further showed that *MALAT1-lo* subpopulations represent early degenerating rod photoreceptors, a finding that has not previously been reported in human or any other species. Previous studies have demonstrated a role of *MALAT1* in regulating the survival of retinal ganglion cells (Li *et al*, 2017) and in pathogenesis of retinal pigment epithelium cells (Yang *et al*, 2016). However, the functional role of *MALAT1* in photoreceptors remained unclear. Our results demonstrated that loss of *MALAT1* expression in rod photoreceptors following putative post-mortem degeneration, and suggests *MALAT1* as a potential target to enhance rod photoreceptor survival and retinal preservation. Future studies are warranted to investigate the functional role of *MALAT1* in photoreceptors. Our transcriptome data also revealed rod photoreceptor clusters specific to particular donor retinas and we showed that application of the CCA method could effectively correct for these donor/batch variations in rod photoreceptors. Further studies with a larger number of donor samples will allow testing of the feasibility of using scRNA-seq to comprehensively analyse the retina in different individuals, such as assessment of the effects of aging or degenerative retinal diseases.

Another outcome of this study is the assessment of biomarkers that allow identification of major retinal cell types and subtypes. Our results provide new insights into the cone photoreceptor subtypes in human. The cone subtypes are traditionally categorized based on expression of different opsins that allowed for cellular detection of light at various wavelength. While the S-cones are structurally different from the other two cone subtypes, the L-cones and M-cones are structurally similar and difficult to distinguish from each other, except for the opsin they expressed (Viets *et al*, 2016). We report the first description of the transcriptome profiles of S-cones in adult human and highlight novel marker genes that can be used to distinguish them. We also identified the transcriptome and novel marker genes for L/M-cones, however given the high sequence homology, particularly at the 3’ end, of *OPN1MW* and *OPN1LW*, we could not confidently separate L-cones and M-cones. In addition, we show that many of the known Müller glia markers are often expressed in retinal astrocytes, and we also provide a detailed assessment of commonly used retinal glial markers showing the differential expression pattern between Müller glia and retinal astrocytes.

Finally, our results highlighted the use of this neural retina transcriptome atlas to benchmark retinal cells derived from stem cells or primary cultures. A major goal of pluripotent stem cell research is to derive cells that are similar to those in adults *in vivo*, which is important for development of stem cell disease models and regenerative medicine (Hung *et al*, 2017). Our analysis shows that hiPSC-derived cone photoreceptors are highly similar to both fetal and adult cones in comparison with all other major retinal cell types. We show that hiPSC-derived cells are more fetal-like than adult-like, which is consistent with other studies (Handel *et al*, 2016; Baxter *et al*, 2015). We also benchmark a commonly used Müller glia cell line MIO-M1. Our results showed that while this cell line exhibits similarities to adult retinal glial cells, there are also some differences between MIO-M1 and adult Müller glia, such as high expression of the glioma-related gene thymosin beta4 (*TMSB4X)* in MIO-M1. Previous reports have also described differences in gene expression in MIO-M1 to Müller glia, and showed that MIO-M1 express markers for post-mitotic retinal neurons and neural stem cells (Lawrence *et al*, 2007; Hollborn *et al*, 2011). These results highlighted the potential effects of prolonged *in vitro* culture of primary retinal cells. Collectively, we showed that the human neural retina transcriptome atlas provides an important benchmarking resource to assess the quality of derived retinal cells, which would have implications for stem cell and neuroscience research.

One of the limitations of this study is the finite number of profiled cell types less frequently represented in the retina such as the amacrine cells and the retinal ganglion cells, which are known to be highly complex. The presented dataset is limited in power to accurately identify differences in the transcriptomes of the subtypes in amacrine and retinal ganglion cells. With the identification of surface markers for these cell types in this study, this work lays the foundation for future research using selection and enrichment (Shekhar *et al*, 2016) of these and other cell types (such as horizontal cells or retinal microglia) to improve the resolution of the human neural retina transcriptome atlas. Another limitation is the use of 3’ gene expression profiling, which presents a challenge for distinguishing L-cones and M-cones. Given the high sequence homology of *OPN1LW* and *OPN1MW* (98%), future studies using full-length mRNA sequencing of single cone photoreceptor cells would provide greater distinction and classification accuracy of the *OPN1MW* and *OPN1LW*-positive cells.

This study describes the transcriptome of human neural retina at a single cell level and is the first to identify the transcriptome of all major human retinal cell types. Our findings shed light on the molecular differences between subpopulations within the rod photoreceptors and the cone photoreceptors. The presented dataset provides an important roadmap to define the genetic signals in major cell types in the human retina and can be used as a benchmark to assess the quality of stem cell-derived cells or primary retinal cells.

## Online Methods

### Human retina collection

Collection of patient samples was approved by the Human Research Ethics committee of the Royal Victorian Eye and Ear Hospital (HREC13/1151H) and Save Sight Institute (16/282) and carried out in accordance with the approved guidelines. For scRNA-seq, post-mortem eye globes were collected by the Lions Eye Donation Service (Royal Victorian Eye and Ear Hospital) for donor cornea transplantation. The remaining eye globes were used for dissection to extract the neural retina. The lens, iris and vitreous were removed and the choroid/RPE layers were excluded from the sample collection. Following extraction, the neural retinal samples were dissociated and processed for scRNA-seq right away. Neural retina samples were dissociated into single cells in dissociation solution (2mg/ml papain, 120 Units/ml DNase I) for 15 minutes. The dissociated neural retina was filtered to ensure single cell suspension using a 30µm MACS Smart Strainer (Miltenyi). For sample from Patient SC, the Dead Cell Removal kit (Miltenyi) was utilised to remove dead cells prior to scRNA-seq. However, in our hands we found that the Dead Cell Removal kit only had a modest improvement in the cell viability (∼8% improvement, data not shown).

### Single cell RNA sequencing (scRNA-seq)

Single cells from three independent neural retina samples were captured in five batches using the 10X Chromium system (10X Genomics). The cells were partitioned into gel beadLinLemulsions and barcoded cDNA libraries, then prepared using the single cell 3’ mRNA kit (V2; 10X Genomics). Single cell libraries were sequenced in 100bp paired-end configuration using an Illumina Hi-Seq 2500 at the Australian Genome Research Facility.

### Bioinformatics processing

The 10X Genomics *cellranger* pipeline (version 2.1.0; (Zheng *et al*, 2017) was used to generate fastq files from raw Illumina BCL files (*mkfastq*) and aligned to the human genome reference GRCh38 using the included STAR alignment software (Dobin *et al*, 2013). Next, *cellranger count* was used to generate read count matrices. To overcome the stringent threshold implemented in cellranger that discards real cells under certain conditions, such as populations of cells with a low RNA content, the *--force-cells* parameter was set to 3000 for the donor 1 library and 5000 for donor 2 and 3 libraries. Using the barcode rank plot, these parameters were selected to increase the number of detected cells for further analysis. The *cellranger* aggregation function (*aggr*) was used to combine the 5 libraries and normalize the between-sample library size differences. The full scRNA-seq dataset is available in ArrayExpress under accession number E-MTAB-7316.

Data were imported into the Seurat single cell analysis software (v2.0.1; https://github.com/satijalab/seurat). Quality control of sequenced libraries was performed to remove outlier cells and genes. Cells with 200-2500 detected genes and expressing < 10% mitochondrial genes were retained. Genes were retained in the data if they were expressed in ≥ 3 cells. Additional cell-cell normalization was performed using the LogNormalize method, and inherent variation caused by mitochondrial gene expression and the number of unique molecular identifiers (UMIs) per cell was regressed out.

Clustering was performed on PCA-reduced expression data using the top 20 principal components using the graph-based shared nearest neighbour method (SNN) which calculates the neighborhood overlap (Jaccard index) between every cell and its nearest neighbors.

Prediction of the cell cycle phase of individual retinal cells was performed in Seurat using cell cycle-specific expression data (Kowalczyk *et al*, 2015). Briefly, genetic markers associated with G2/M and S phase were used to assign cell scores, and cells expressing neither of the G2/M or S phase markers were classified as being in G1 phase.

Sequencing data for fetal (scRNA-seq) and hiPSC-derived cone photoreceptors (bulk RNA-seq) was obtained from ArrayExpress using the accession numbers E-MTAB-6057 and E-MTAB-6058 (Welby *et al*, 2017). Gene expression matrices were generated from the fastq files using the STAR aligner software. scRNA-seq data from 72 cells were quality-controlled, filtered and then normalised with the scran algorithm (Lun *et al*, 2016) as described (Welby *et al*, 2017), using the *ascend* (https://github.com/IMB-Computational-Genomics-Lab/ascend) package in R, which resulted in 63 high quality single cell transcriptomes. Bulk RNA-seq data generated from 6 hiPSC-derived cone photoreceptor cultures was filtered such that each gene was represented by at least 10 counts in all samples and normalisation was performed in edgeR using the trimmed mean of M method (Robinson & Oshlack, 2010). Pre-processed scRNA-seq data generated from adult retina (Phillips *et al*, 2018) was obtained from the Gene Expression Omnibus (GSE98556).

### Canonical correlation analysis

Canonical correlation analysis (CCA) was applied to correct donor-specific effects observed in the rod photoreceptor populations. This was achieved by separating the raw data into 5 sample-specific datasets, which were then used as inputs for the RunMultiCCA function in Seurat. For the CCA analysis, we used the most highly variable genes that were shared by all 5 samples and the recombined data was aligned using the first 20 CC dimensions, selected by biweight midcorrelation (bicor) analysis. Aligned cells were reclustered in Seurat using the first 20 aligned CC dimensions at a resolution of 0.6

### Identification of retinal cell types

Cell types were classified using differential expression analysis, which compared each cluster to all others combined using the Wilcoxon method in Seurat to identify cluster-specific marker genes. Each retained marker gene was expressed in a minimum of 25% of cells and at a minimum log_2_ fold change threshold of 0.25.

In paired cluster analyses, differentially expressed genes were considered significant if the adjusted p-value was less than 0.01 (Benjamini-Hochberg correction for multiple testing) and the absolute log_2_ expression fold change was ≥ 0.5.

### Mapping cells between subpopulations in different samples

To compare subpopulations identified in the merged dataset (5 samples), we applied scGPS (single cell Global Projection between Subpopulations), a machine learning procedure to select optimal gene predictors and to build prediction models that can estimate between-subpopulation transition scores. The transition scores are the probability of cells from one subpopulation that are in the same class with the cells in the other subpopulation. The scores, therefore, estimate the similarity between any two subpopulations. Here, we compared three main subpopulations from sample Retina 2A with all subpopulations in the sample Retina 2B. The source code of the scGPS method is available with open-access (https://github.com/IMB-Computational-Genomics-Lab/scGPS).

### Correlation of scRNA-seq data with retinal cell types

The mean expression levels of cells in each cluster were calculated and used to calculate Pearson’s correlations in a pairwise manner with each of the other clusters and results were deemed significant if the correlation P-value was less than 0.01.

### Pathway analysis

Gene enrichment analysis was performed using Enrichr (Kuleshov *et al*, 2016). The combined score, computed by taking the log of p-value from the Fisher exact test and multiplying by the z-score of the deviation of the expected rank, was used to determine the enrichment ranking for pathways, othologies, transcription factor network and protein network analysis.

### Fluorescent in situ hybridization

Donor retinas were first dissected from the eye cup. The retinal tissues were subjected to 30% sucrose cryoprotection and were then frozen in -80°C. Sections were cut on a cryostat (Leica CM3050S) and mounted on glass slides (SuperFrostPlus). The retinal samples were fixed in 3.7% (vol/vol) formaldehyde and hybridized with Stellaris RNA FISH probes (Biosearch Technologies) against *MALAT1* labeled with Quasar 570, following the manufacturer’s instructions. Briefly, samples were incubated with Quasar 570-labeled probes at 125nM in hybridization buffer and hybridized 5 hours at 37°C, followed by nuclear counterstain using DAPI. The samples are imaged using a ZEISS confocal laser-scanning microscope (ZEISS, LSM700).

## Author Contributions

HW, AM coordinated and collected the human donor retinas, GP provided the funding for human donor tissue collection, LF, TN, JJ, SH, RW, LZ, TZ conducted the experiments; SL, CL, AS, RW, QN, EW, JS, TL, LZ, TZ processed and/or analysed the data; PYW, AH, JP, MG and RW contributed to experimental design and data analysis; SL, AH, JP and RW wrote the manuscript.

## Acknowledgements

The human Müller cell line Moorfields/Institute of Ophthalmology-Müller 1 (MIO-M1) was obtained from the UCL Institute of Ophthalmology, London, UK. The authors want to thank David Mackey and Holly Chinnery for critical discussion and helpful feedback for this study, and Prema Finn for technical assistance for the collection of human donor samples. This work was supported by funding from the Ophthalmic Research Institute of Australia (RW, SL), the University of Melbourne De Brettville Trust (RW) and the Kel and Rosie Day Foundation (RW). JCS is supported by GOSHCC. The Centre for Eye Research Australia receives operational infrastructure support from the Victorian Government.

## Supplementary figures and tables

**Supplementary table 1:** Details for donor retina samples.

| Retina | Patient ID | Eye bulb | Sex | Age (years) | Retrieval time (hrs) | Ocular complications | Assay | scRNA-seq Library | Targeted cell number | Captured cell number after QC |
| --- | --- | --- | --- | --- | --- | --- | --- | --- | --- | --- |
| 1 | 17-010 | Right | F | 80 | 11.5 | Cataract on left eye | scRNAseq, cell viability | A | 4000 | 2122 |
| 2 | SC | Left | M | 42 | 6.2 | - | scRNAseq, cell viability | A | 10000 | 4449 |
|  |  |  |  |  |  |  |  | B | 10000 | 4528 |
| 3 | 17-011 | Right | F | 53 | 14.5 | - | scRNAseq, cell viability | A | 10000 | 4518 |
|  |  |  |  |  |  |  |  | B | 10000 | 4392 |
| 4 | 16033 | Left | M | 73 | 12 | - | FISH |  |  |  |
| 5 | 16061 | Left | M | 79 | 25 | - | FISH |  |  |  |
| 6 | 16088 | Left | F | 61 | 22 | - | FISH |  |  |  |
| 7 | 16168 | Left | M | 74 | 7, 13 | - | FISH |  |  |  |
| 8 | 17-080 | Right | F | 77 | 4.25 | - | Cell viability |  |  |  |
| 9 | 17-143 | Left | M | 66 | 23 | - | Cell viability |  |  |  |
| 10 | 17-153 | Left | M | 73 | 13 | - | Cell viability |  |  |  |
| 11 | 17-160 | Left | F | 76 | 33.75 | - | Cell viability |  |  |  |
| 12 | 17-167 | Right | M | 49 | 16 | - | Cell viability |  |  |  |

**Supplementary table 2:** Breakdown of cell number assigned to individual donor libraries. The red highlight the number of cells assigned in the largest clusters, showing similar cell assignment to clusters between 2 replicates.

| Clusters | Retina 1 | Retina 2A | Retina 2B | Retina 3A | Retina 3B |
| --- | --- | --- | --- | --- | --- |
| 0 | 58 | 1927 | 1924 | 103 | 66 |
| 1 | 307 | 33 | 30 | 1419 | 1330 |
| 2 | 3 | 1275 | 1299 | 9 | 9 |
| 3 | 172 | 413 | 425 | 447 | 670 |
| 4 | 74 | 23 | 46 | 999 | 864 |
| 5 | 232 | 59 | 74 | 398 | 384 |
| 6 | 156 | 170 | 139 | 273 | 264 |
| 7 | 737 | 6 | 8 | 42 | 41 |
| 8 | 97 | 98 | 124 | 208 | 190 |
| 9 | 106 | 184 | 189 | 93 | 86 |
| 10 | 33 | 83 | 80 | 180 | 188 |
| 11 | 42 | 83 | 68 | 93 | 77 |
| 12 | 55 | 35 | 35 | 108 | 82 |
| 13 | 18 | 20 | 40 | 58 | 51 |
| 14 | 14 | 7 | 11 | 62 | 71 |
| 15 | 11 | 4 | 12 | 13 | 9 |
| 16 | 5 | 8 | 11 | 9 | 9 |
| 17 | 2 | 21 | 13 | 4 | 1 |

**Supplementary table 3:** Membrane-related markers were extracted from GO annotation using the differentially expressed genes in the major human retinal cell types.

| **Cell types** | **Identified surface markers** |
| --- | --- |
| Rod photoreceptors | *RHO, ROM1, CNGA1, CNGB1, PRPH2, MFGE8* |
| Cone photoreceptors | *OPN1LW, SLC24A2, DST, TTR* |
| Muller glia | *RGR, GPR37, RLBP1, NCAM1, CD164* |
| Horizontal cells | *CNTNAP2, PTN, NDRG4, TAGLN3* |
| Bipolar cells | *TRPM1, GRM6* |
| Amacrine cells | *EPHB6, SLC5A7, GABRD, KCNJ12, PTPRF, GRIN2A* |
| Retinal astrocytes | *GYPC, SERPINA5, CD44, SLC4A4, AQP4, ABCC3, NGFR* |
| Microglia | *CD74, IILA-DRA, IILA-DPA1, TYROBP, FCER1G, LAPTMS, CD14, HLA-DQA1, FXYD5, CTSB, CD37* |
| Retinal ganglion cells | *RTN1, NDRG4, UCHL1, YWHAH* |

**Supplementary figure 1:**
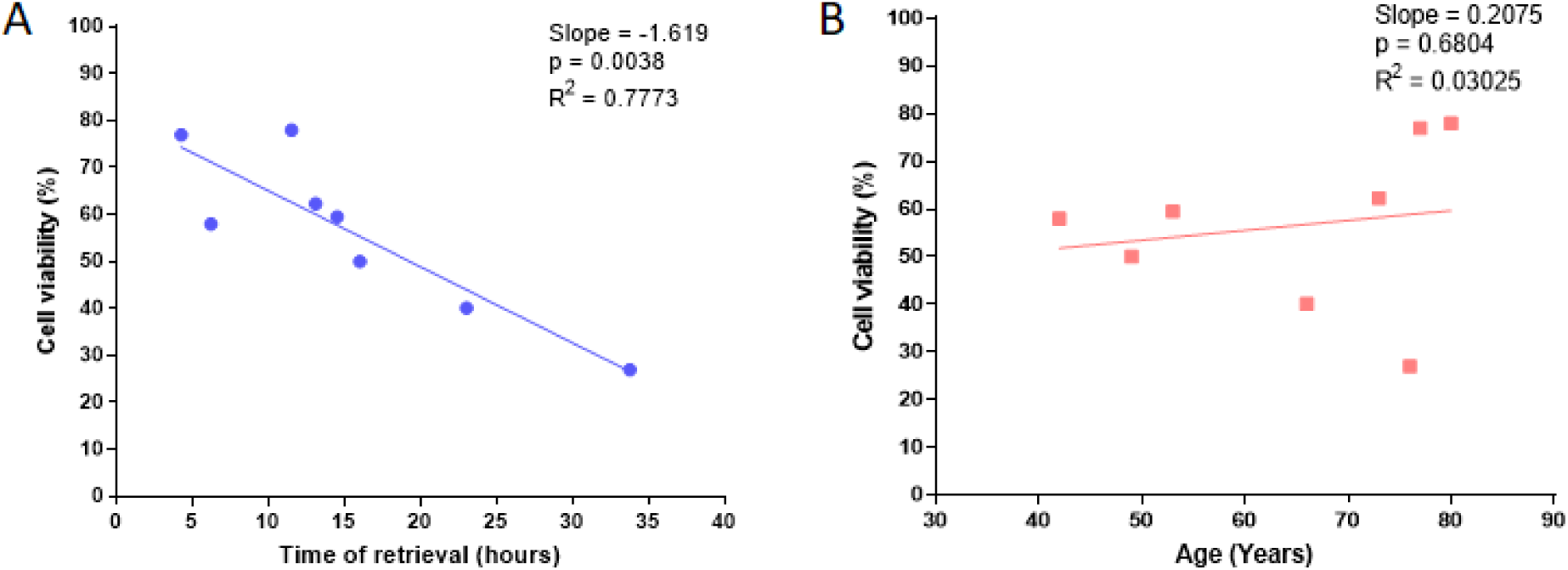
Correlation of cell viability of post-mortem human neural retina with A) time of tissue retrieval after death, and B) age of donor.

**Supplementary figure 2:**
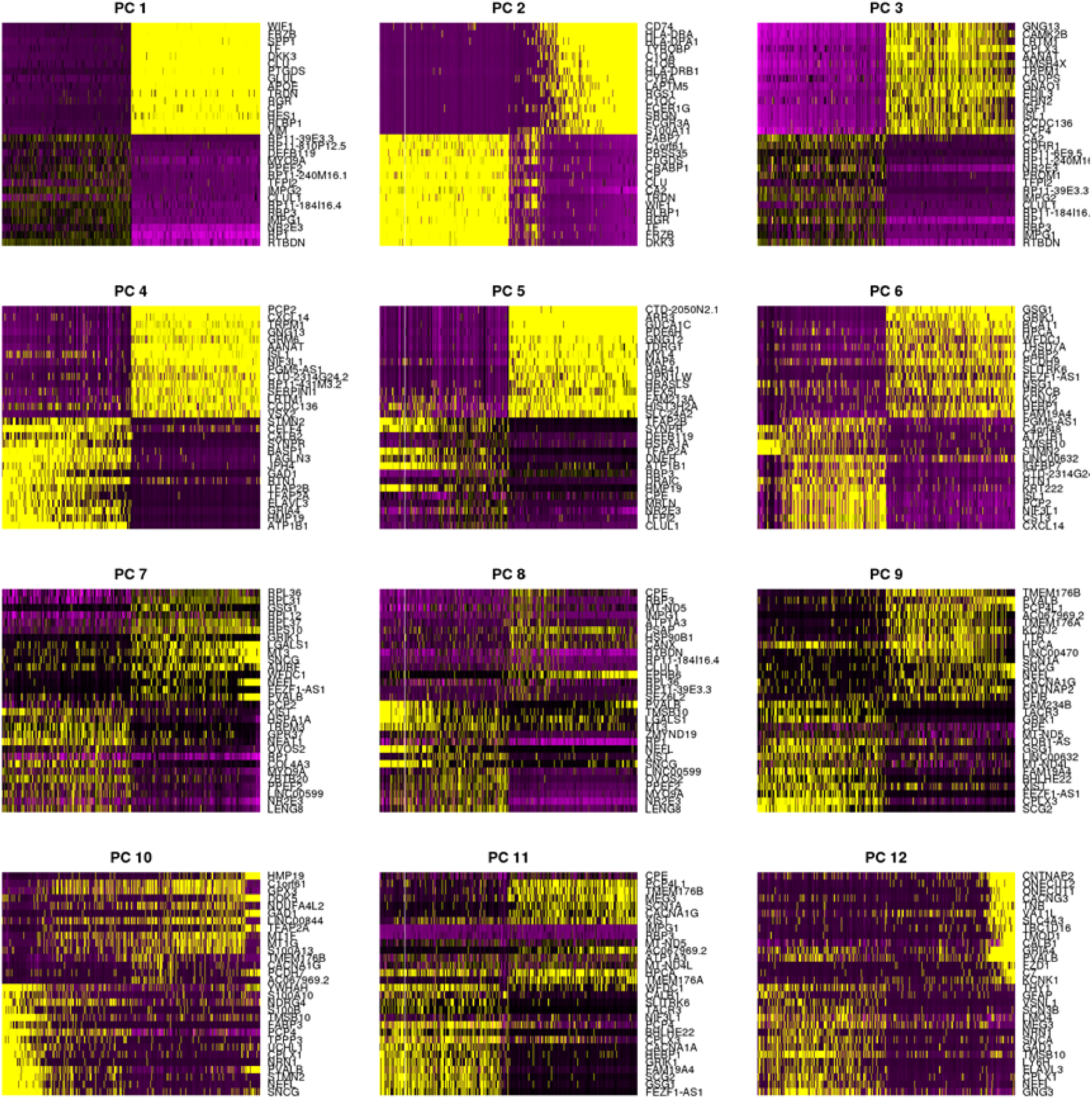
Heatmaps of the top 12 principal components explaining the primary sources of heterogeneity in the retinal scRNA-seq data. Cells and genes are ordered by PCA score calculated by Seurat. The genes driving the majority of the variance are determined using the top 500 cells.

**Supplementary figure 3:**
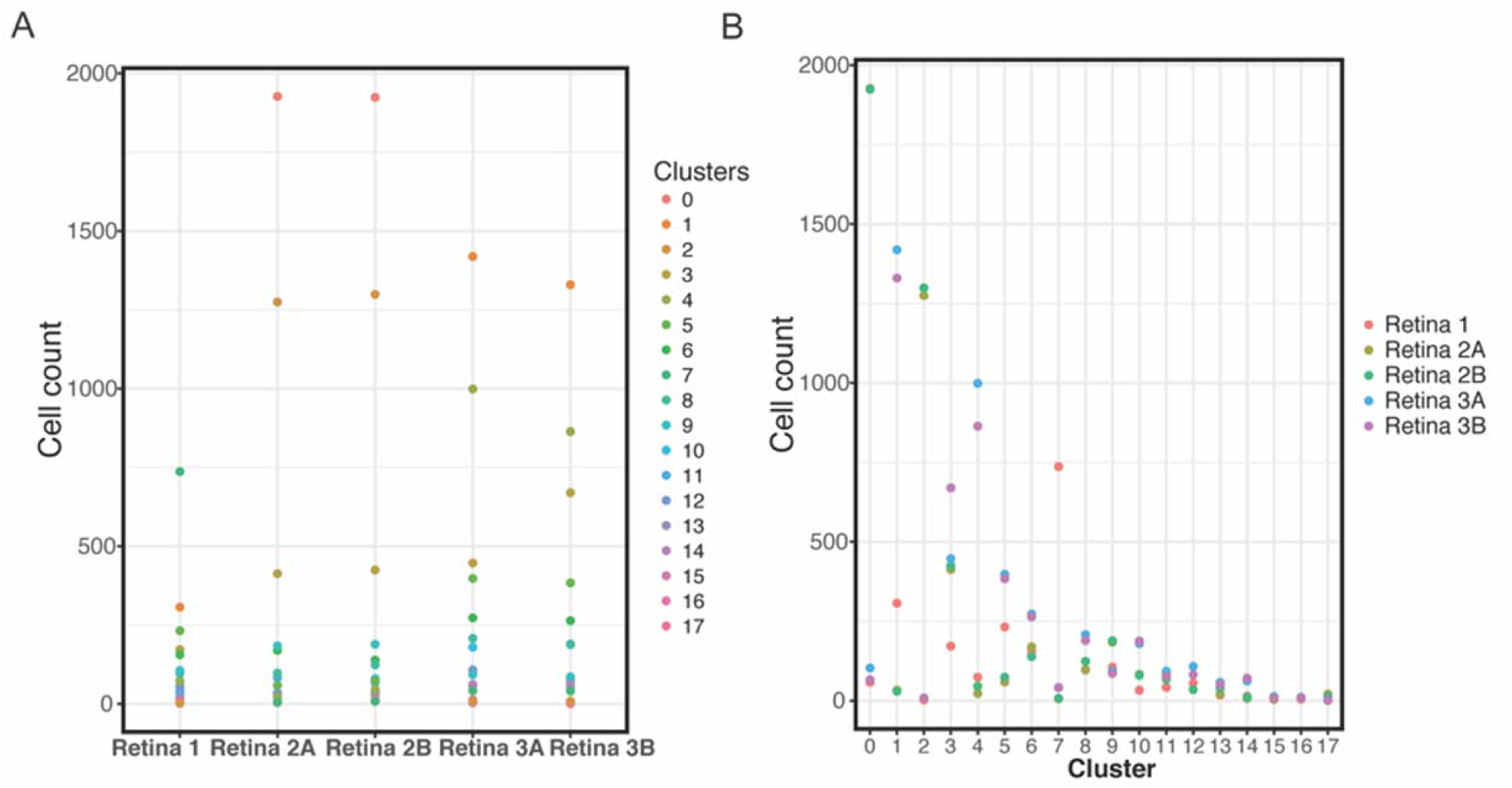
Distribution of frequency of the 18 clusters (C0-C17) in individual single cell libraries ordered by A) individual single cell library or B) identified clusters.

**Supplementary figure 4:**
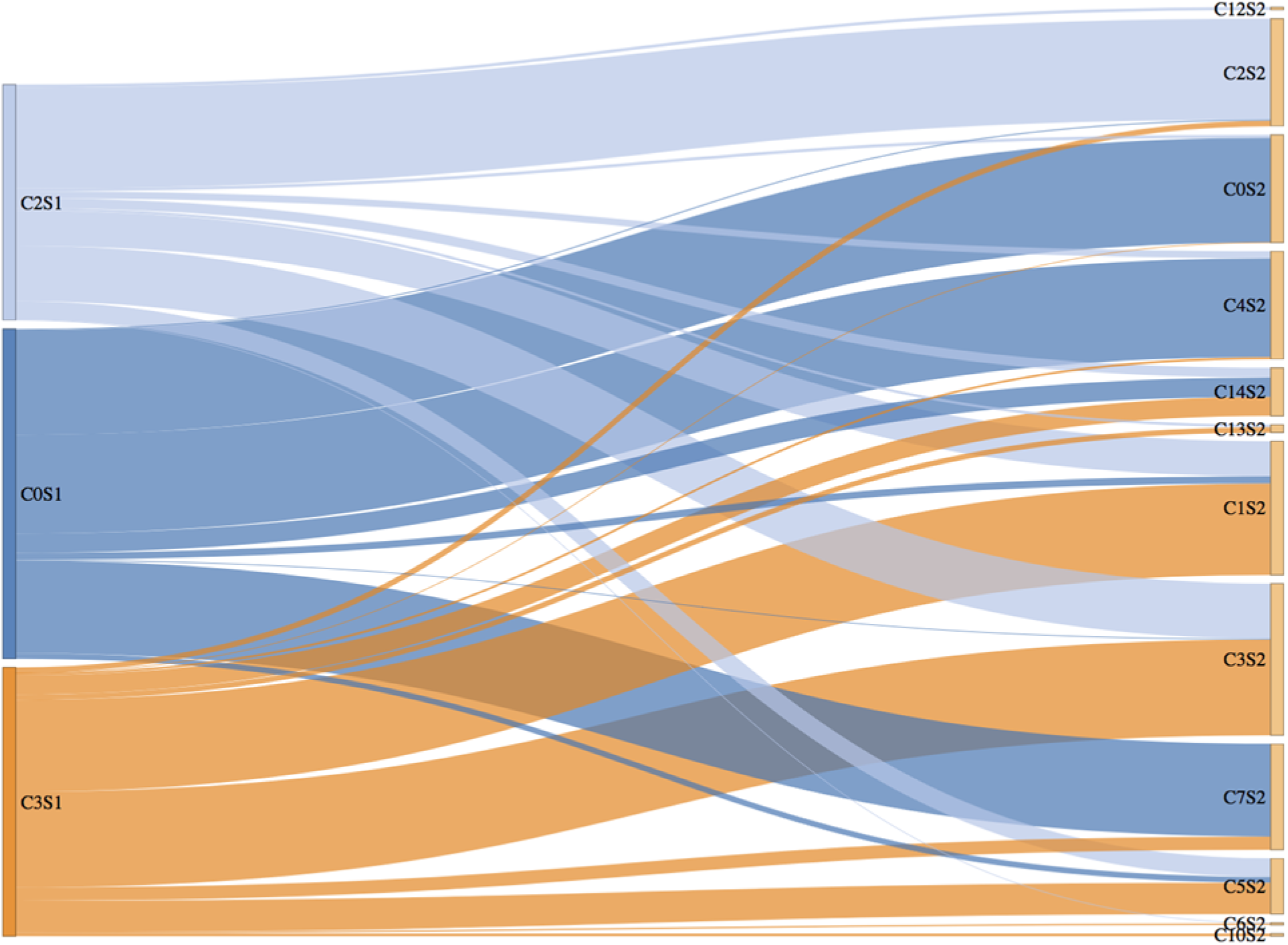
Prediction of cluster relationship between library technical replicates in Retina 1 and Retina 2. The Sankey plot shows edges connecting clusters, with larger edge indicating higher similarity, ranging from 0 to 100%. The size of the edge was quantitatively estimated by implementing scGPS modelling approach for pairs of clusters, as described in the method section. The three largest clusters in Retina 2A were compared with all clusters in Retina 2B. Consistently we see C2 in Retina 2A is most similar to C2 in Retina 2B. The same trend is seen for C0 and C3. These results demonstrated that the variation between library replicates is minimal in our dataset, and that the clusters determined from the merged dataset were consistent across samples. We also found higher similarities among Rod photoreceptor clusters (C0, 2, 3 in Retina 2A with clusters C0, 2, 4, 7 in Retina 2B) than compared with other clusters.

**Supplementary figure 5:**
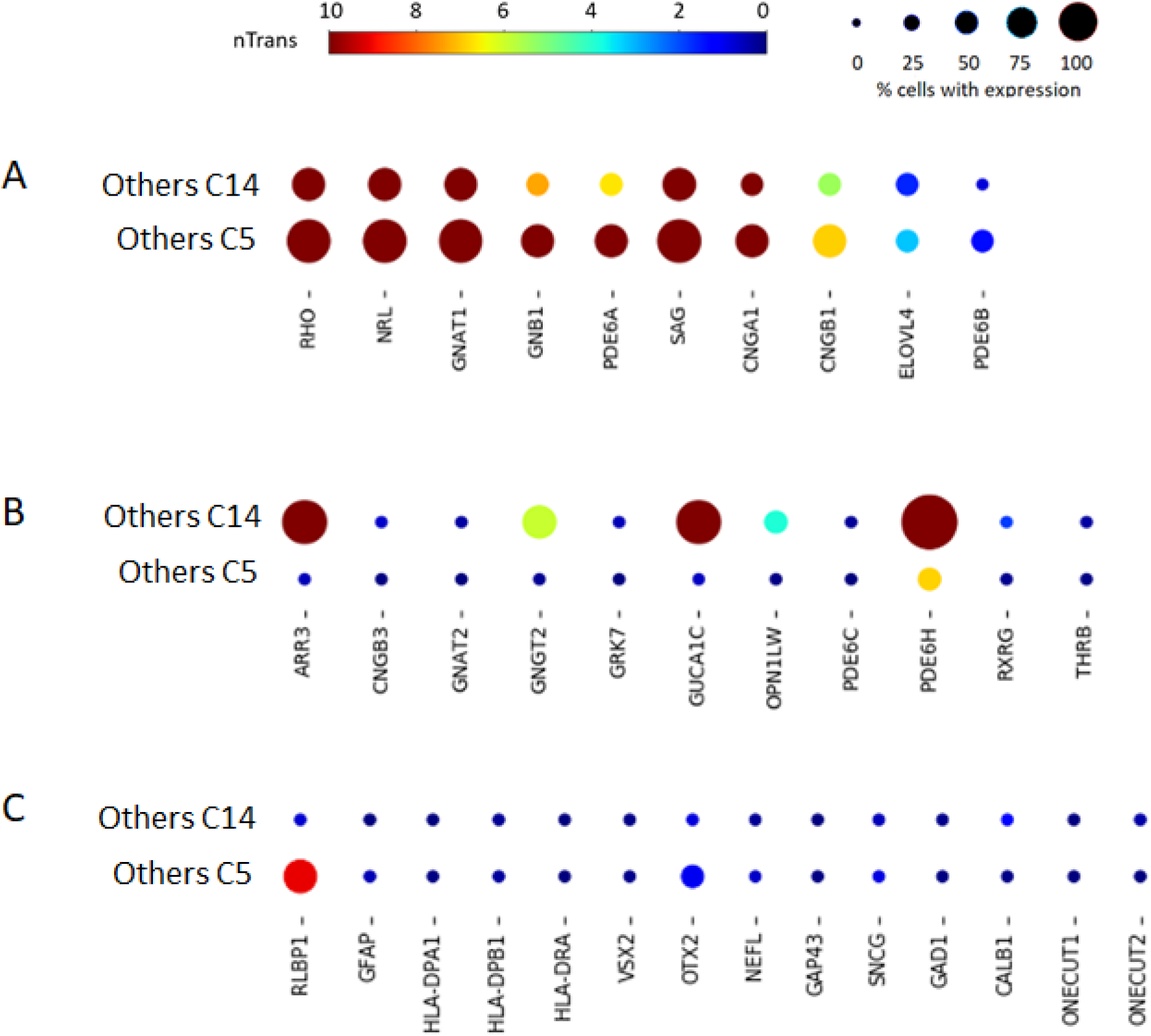
Feature expression heatmap showing expression patterns of A) rod photoreceptor markers, B) cone photoreceptor markers and C) other major retinal class markers (Müller glia: RLBP1; astrocytes: GFAP; microglia: HLA-DPA1, HLA-DPB1, HLA-DRA; Bipolar cells: VSX2, OTX2; retinal ganglion cells: NEFL, GAP43, SNCG; Amacrine cells: GAD1, CALB1; Horizontal cells: ONECUT1, ONECUT2) in unassigned clusters C14 and C5. The size of each circle depicts the percentage of cells expressing the marker within the cluster.

**Supplementary figure 6:**
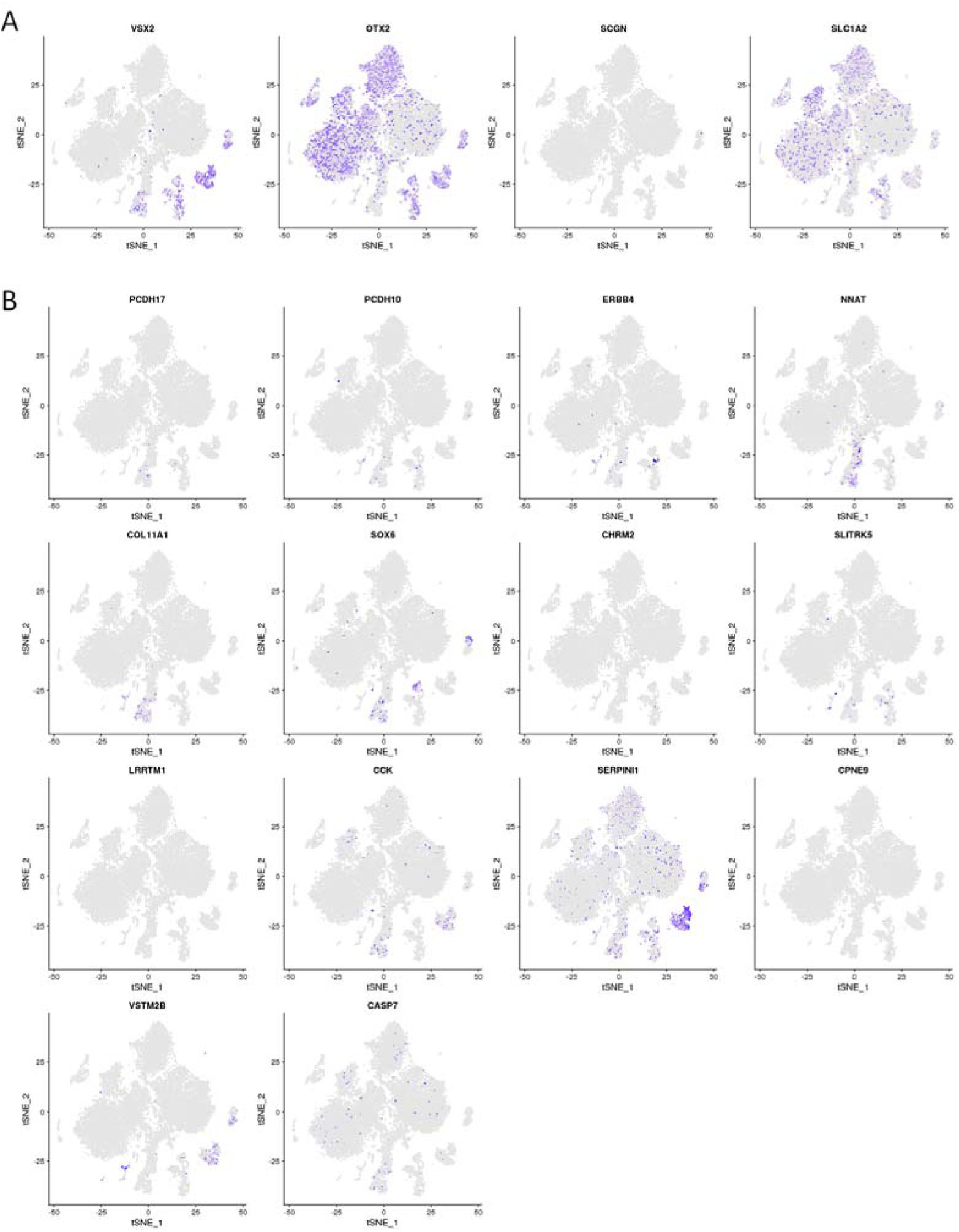
t-SNE plots showing gene expression in the compiled human neural retina transcriptome atlas (20,009 cells) for A) 4 commonly used bipolar markers and B) 14 new markers for individual bipolar subtypes identified in previous mouse scRNA-seq study (Shekhar *et al*, 2016).

**Supplementary figure 7:**
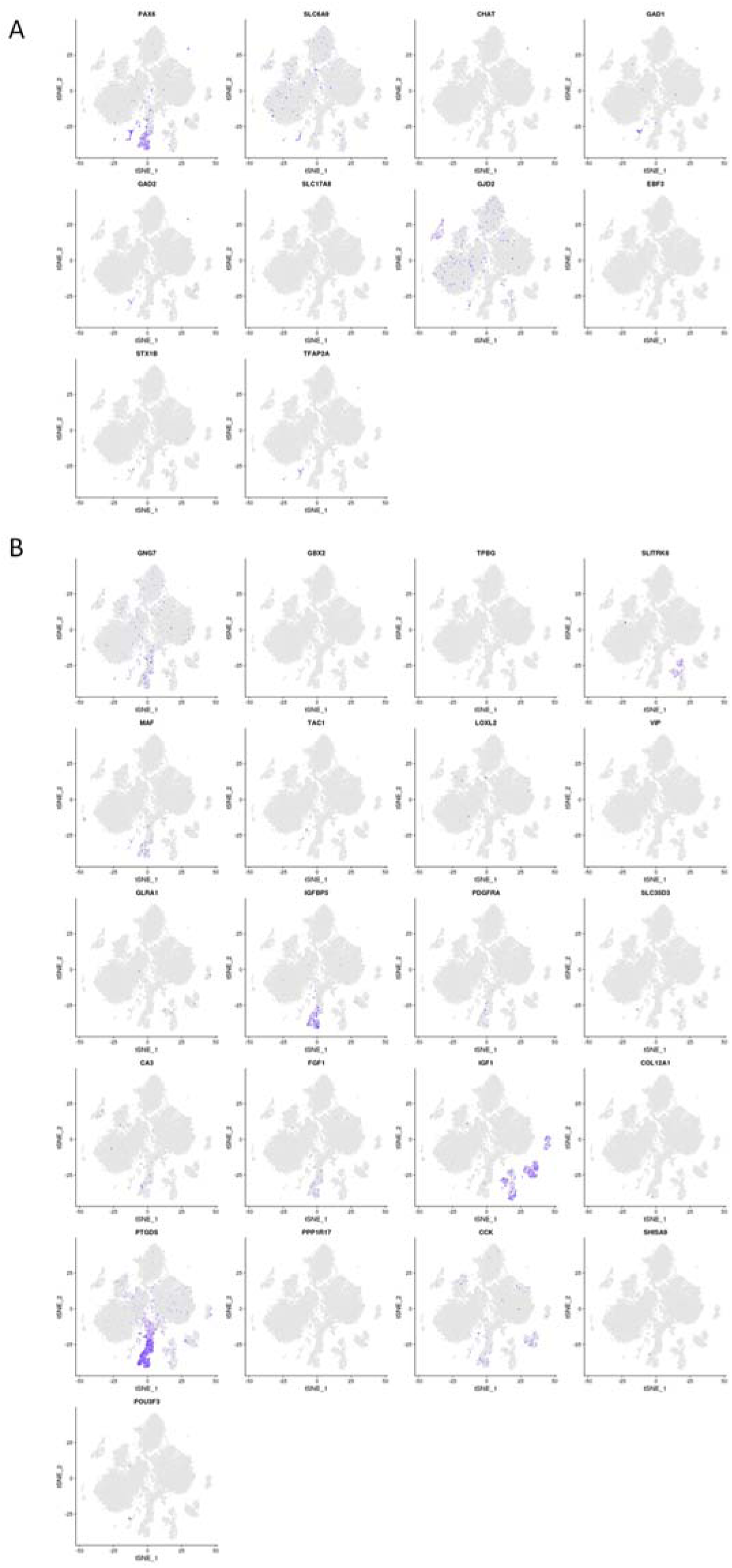
t-SNE plots showing gene expression in the compiled human retina transcriptome atlas (20,009 cells) for A) 10 commonly used amacrine markers and B) new markers for amacrine subtypes identified in previous mouse scRNA-seq study (Macosko *et al*, 2015).

**Supplementary figure 8:**
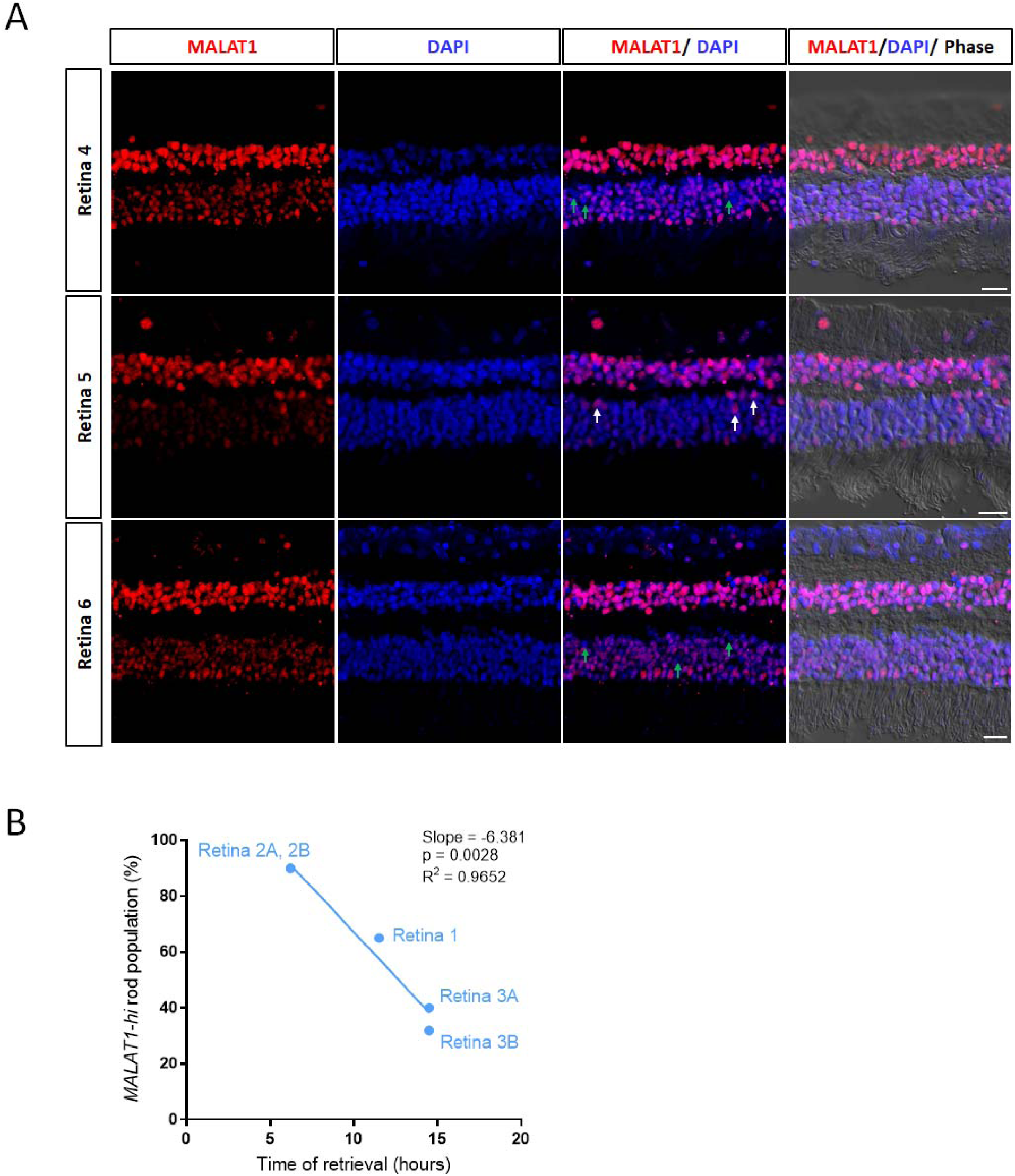
Fluorescent in situ hybridization showing expression of *MALAT1* in three donor retina samples (Retina 4-6). Green arrows highlight rod photoreceptors with low levels of *MALAT1* in Retina 4 and 6, white arrows highlight rod photoreceptors with high levels of *MALAT1* in Retina 5. Scale bars = 20µm. B) Correlation of proportion of MALAT1-hi rod populations with time of retina retrieval after death for Retina 1-3.

**Supplementary figure 9:**
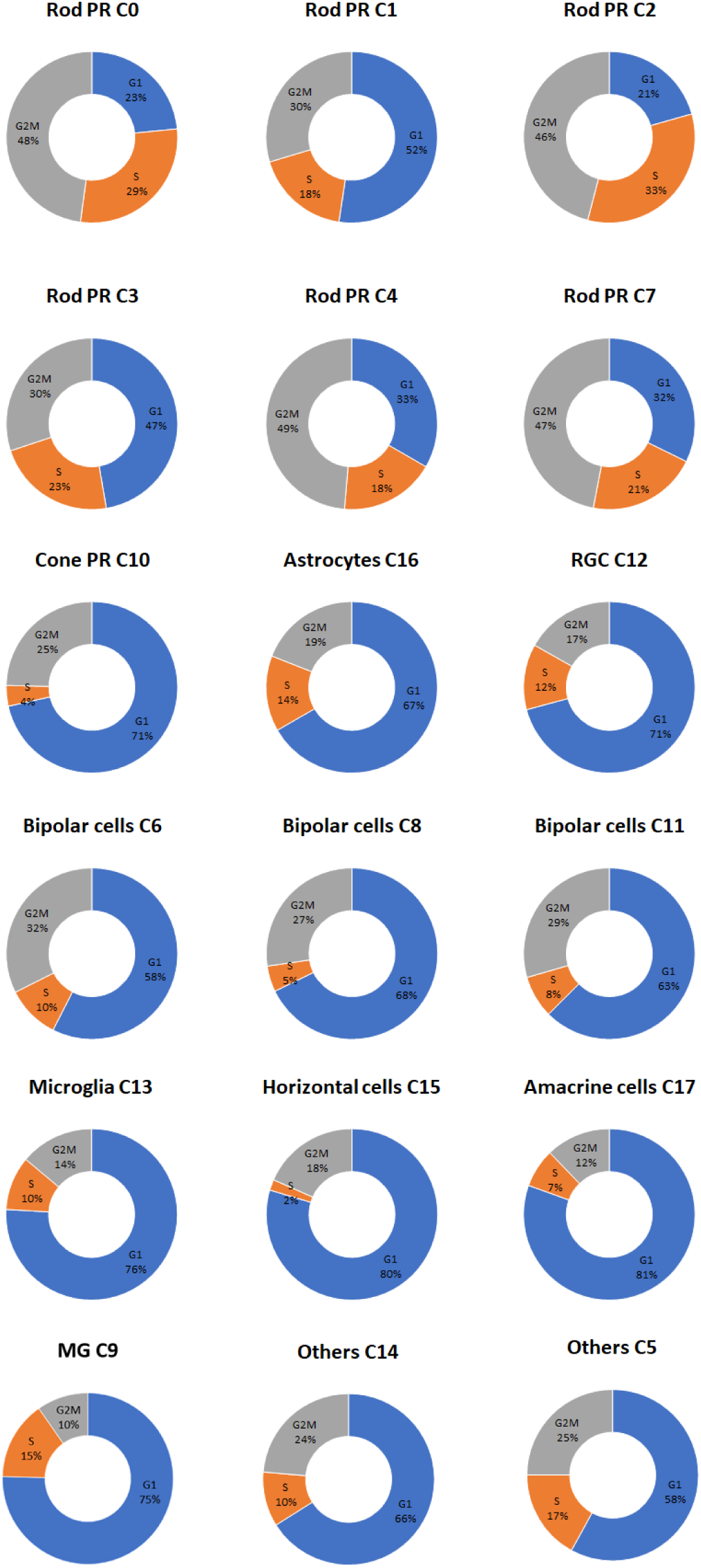
Cell cycle scores across major retinal cell clusters showing the likelihood for the proportion of cells in G1, S or G2/M phases.

**Supplementary figure 10:**
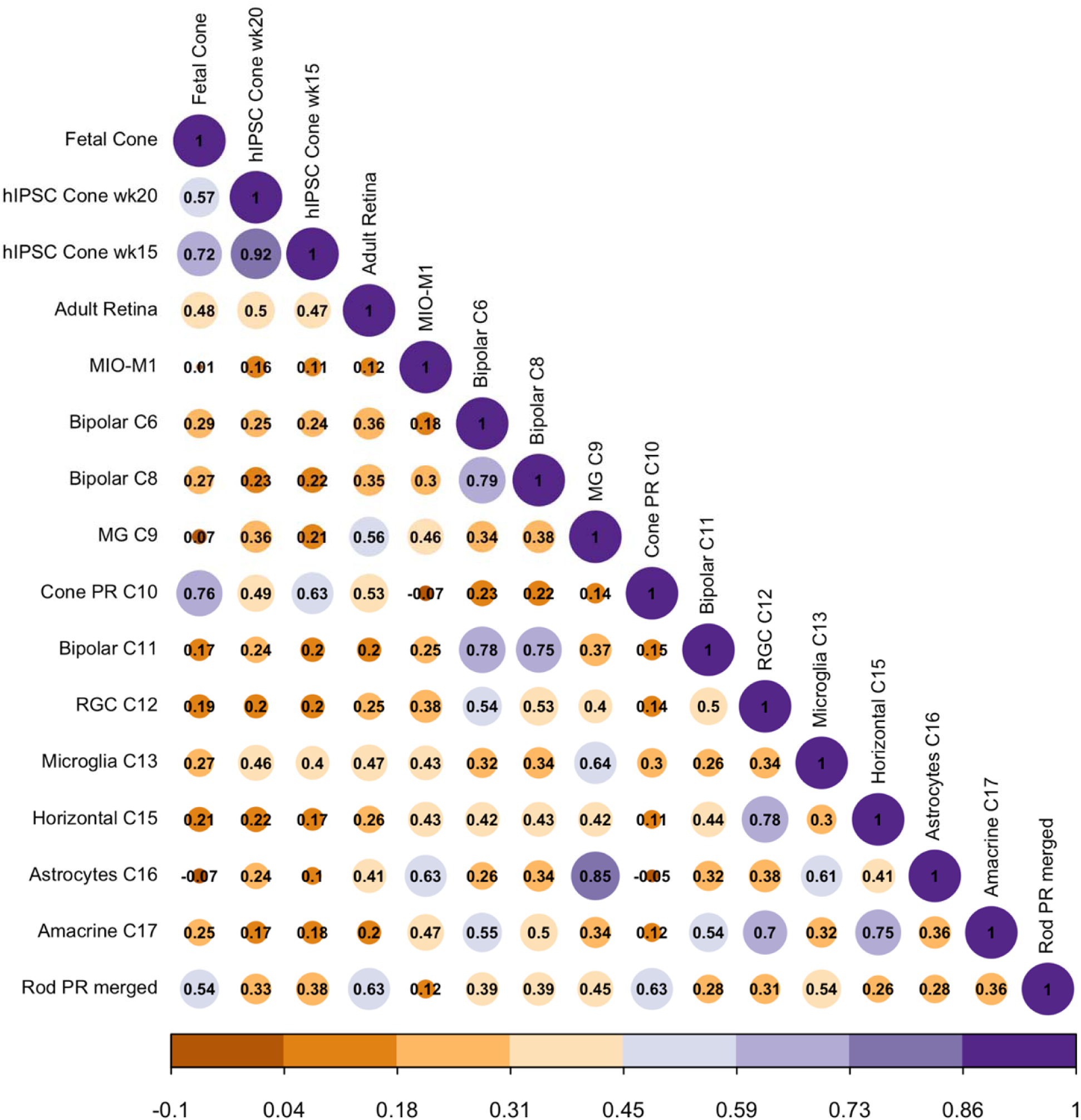
Correlation matrix to benchmark hiPSC-derived cone photoreceptors (week 15, week 20; (Welby *et al*, 2017), fetal cone photoreceptors (Welby *et al*, 2017), adult retina (Phillips *et al*, 2018) and the human Müller glia cell line MIO-M1 against all retinal cell types identified in this human neural retina atlas.

